# Eavesdropping on Herbivores: Using contact microphones to quantify Plant-Insect Interactions

**DOI:** 10.1101/2024.09.23.614472

**Authors:** Dev Mehrotra, Laurence Still, Vidit Agrawal, Kimberly Gibson, James D. Crall, Emily N. Bick

**Affiliations:** Department of Computer Science, University of Wisconsin-Madison, WI, USA; Department of Entomology, University of Wisconsin-Madison, WI, USA

**Keywords:** contact microphones, herbivorous insects, pest monitoring, acoustic monitoring, spectrogram analysis

## Abstract

1. Herbivorous insects are major crop pests whose feeding often results in significant economic damage. Detecting and monitoring insect herbivory is crucial for effective pest management, but manual methods can be time-consuming, inefficient, and destructive. This study investigates the use of contact microphones as a non-invasive and efficient tool for detecting, identifying, and monitoring insects feeding on plants.
2. Approach and methods: Contact microphones connected to a Raspberry Pi were clipped onto plant stems and programmed to record sounds made by the feeding insects. Three combinations of herbivorous insects and plants were evaluated: tobacco hornworm (*Manduca sexta* Linnaeus) on tobacco, Colorado potato beetle (*Leptinotarsa decemlineata* Say) on potato, and European corn borer (*Ostrinia nubilalis* Hübner) and Northern corn rootworm (*Diabrotica barberi* Smith & Lawrence) on corn (*Zea mays* Linnaeus). Sound recordings were analyzed to determine the presence and absence of insects and to quantify differences in feeding activity between different insects. The laboratory study was repeated in a corn field to observe corn rootworms (*Diabrotica* spp.).
3. Main results: Contact microphones successfully detected insect herbivory on the tobacco, potato, and corn plants in the lab and field, demonstrating the use of substrate-born acoustic signals for detecting both large and small-bodied insect pests. The recordings also revealed differences in the feeding patterns, frequency, and amplitude between insect species. Field experiments indicate low background noise relative to lab experiments.
4. Conclusions and implications: Contact microphones offer a promising cost-effective, non-destructive, and efficient method for monitoring insect herbivory. This approach has the potential to improve scientific observations of insect herbivores and pest management strategies by enabling early pest detection and measuring the success of interventions. Early field results indicate the viability of this approach under realistic conditions in the field. The results highlight the importance of considering the acoustic environment in agricultural ecosystems, suggesting sound can be used to better understand the behavior of insects and their interactions with plant hosts. Contact microphones could have widespread applications in agriculture and ecology by providing a simple and effective tool for detecting and monitoring insect activity.

## 1. Introduction

Herbivorous insects pose a significant threat to global food security and are responsible for approximately∼US$470 billion in annual economic losses (Sharma et al., 2017). Insect pest damage results in an estimated 20% reduction in yield globally (Godfray et al., 2010), with climate change predicted to exacerbate these phenomena (Lehmann et al., 2020).

Time efficiency and labor costs present notable challenges for monitoring herbivorous insects in agroecosystems (Subramanian et al., 2021), often resulting in limited or outdated pest data and therefore increasing uncertainty in decisions (Shea et al., 2002). Moreover, inadequate monitoring may result in excessive use of pesticides (Pomari-Fernandes et al., 2015), which have unintended negative environmental effects (Rezác et al., 2019), such as negative impacts on non-target, beneficial insects, and can lead to suboptimal pest mitigation (Coslor et al., 2019; Gagic et al., 2021). Conventional monitoring procedures are often destructive to crops, particularly when observing insects that feed within plant tissues (i.e. boring insects) (Mcguire et al., 1957) or on roots (Gassmann et al., 2012). Degree-day models that use accumulated temperature to predict insect development timing offer a more precise approach to timing interventions (Bick et al., 2020). However, insects’ rapid evolution in response to abiotic conditions such as insecticides (Chen et al., 2023) and the changing climate (Garnas, 2018) can render these models inaccurate. Therefore, new tools are needed to study and inform the mitigation of these rising herbivore threats (Preti et al., 2021; Zijlstra et al., 2011).

Automation of insect monitoring provides an opportunity to overcome the challenges of conventional monitoring and improve management decisions (Hagstrum et al., 1996; Kirkeby et al., 2021; Pegoraro et al., 2020; Potamitis et al., 2017; Rydhmer et al., 2021). Automated monitoring methods for identifying insects are diverse, including identification in flight with optical sensors (Kirkeby et al., 2021; Rydhmer et al., 2021) and audio sensors (Hassall et al., 2021), within traps (Blair et al., 2020; Preti et al., 2021) or on plants with cameras (Bjerge Id et al., 2023), from photos (Wäldchen & Mäder, 2018), and underwater from acoustic recordings (Desjonquères et al., 2020). High sensor costs, however, often limit scalability for both scientific observation and practitioner adoption. Additionally, with limited exceptions (Rach et al., 2013), existing methods are not suitable for observing insects that are underground or feeding within plant tissues.

To address cost constraints and broaden the observational scope of insect sensors, here we demonstrate the application of clip-on piezo-electric microphones as an affordable, non-destructive tool for the detection, identification, and monitoring of insect herbivory on plants – via eavesdropping. The application of piezo-electric microphones to automate insect monitoring could lead to scalable observations with use cases in biology, ecology, agriculture, and beyond. Therefore, we hypothesized that a contact microphone clipped onto a plant enables detection of insect herbivory. This hypothesis was evaluated by examining four species belonging to two highly diverse orders (Coleoptera and Lepidoptera), encompassing two large-bodied and two small-bodied insect species in the lab, and one root-feeding species in the field.

## 2. Materials and Methods

### 2.1 Hardware

The insect sensor (hereafter referred to as ‘the sensor’) is composed of a Raspberry Pi 4 (4GB RAM), an optional USB hub (H302S USB 3.0 Hub, SmartQ, Taipei, Taiwan), one to four clip-on contact microphones (Clip On Contact Microphone Piezo Pickup 3m 6.35mm, Alomejor Store, China), an adapter (1/4’’ to 3.5mm Stereo Pure Copper Headphone Adapter, 3.5mm (1/8’’) Plug Male to 6.35mm (1/4’’) Jack Female Stereo Adapter, ANDTOBO, Shenzhen, Guangdong, China), a USB sound card (USB External Stereo Sound Adapter, SABRENT, Los Angeles, CA, USA), a 5 volt power source (5 Volt AC adapter, CanaKit, North Vancouver, BC, Canada), and a memory card (Evo 128 GB, Samsung, Seocho-gu, South Korea) (Table 1, as described in US Provisional Patent #63/463,350). The sensor records sound (i.e., insect chewing, and tapping) from the contact microphone component clipped onto the plant. Each Raspberry Pi can host up to four microphones for continuous observation. The components are assembled using the plug and play function on the Raspberry Pi (see Figure 1).

**Table 1.** A table of components, prices (at time of experiment), and sources.

| **Part** | **Cost** | **Purchased from** |
| --- | --- | --- |
| **Cana Kit Raspberry Pi 4 Extreme (4GB RAM)** | \$169.99 | Amazon Canakit |
| **Contact microphones** | \$7.97 | Amazon Amity EXP |
| **6.35mm male to 3.55m female adapter** | \$8.95 | Amazon ANDTODO-US |
| **TYPE C Cable and Power Adapter** | Included with Raspberry Pi | Amazon Canakit |
| **128 GB memory card** | Included with Raspberry Pi | Amazon Canakit |
| **USB Sound Card** | \$7.58 | Amazon Store4PC |

**Figure 1.**
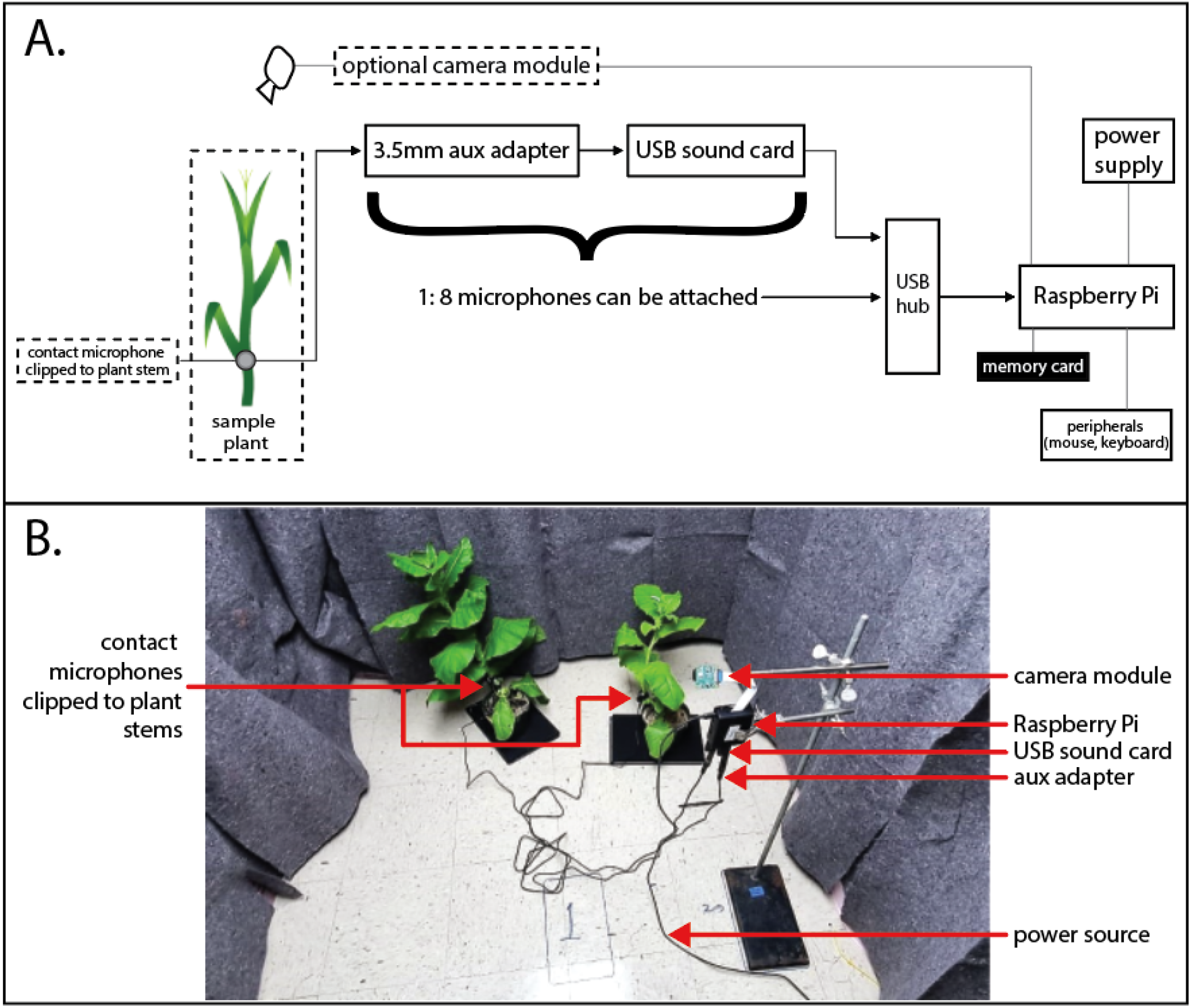
**A**. Diagram of sensor components. **B**. Photo of the assembled sensor device recording sound and video from a laboratory experiment consisting of two tobacco plants, one with and one without a tobacco hornworm.

### 2.2 Software

Recording software enables the sensor to log and export sound gathered by the contact microphone or microphones. The recording software was written in Python using the ‘sounddevice’ package (Geier, 2023). The source code for the sensor’s recording software is stored in a GitHub repository (https://github.com/wiscbicklab/InsectEavesdropperMethodsPaper).

To record sound, a thread is initiated for each microphone with a preset recording duration. Manipulatable variables for the sound recording are the recording duration (preset at one hour), the sampling rate (44,100 Hz), and the file name. The program iteratively saves the data from each microphone as a ‘.wav’ file with the file name and a microphone ID to the memory card while running. Once all open threads are finished the program checks the completion status. The file is accessed by removing the memory card from the Raspberry Pi and transferring it to a standard computer. The data was processed by extracting peaks above the baseline background noise. This ‘baseline’ is defined as the signal vibrations present in the surrounding environment. This noise is highly consistent throughout the entire sound file. The peaks extracted represent insect interactions with the plants. Moreover, the amplitude feature on the signal was extracted using ‘Librosa’ library in Python 3.10.12.

### 2.3 Laboratory experiment

The sensor’s capacity to identify acoustic signals from insect herbivory was evaluated on four insect species within Lepidoptera and Coleoptera. As size is likely to significantly affect acoustic signals, the sensor was tested against two large-bodied leaf-feeding insects and two small-bodied insects, one stem boring, and the other root-feeding. The large-bodied insects were tobacco hornworm (*Manduca sexta* (Lepidoptera: Sphingidae)) and Colorado potato beetle (*Leptinotarsa decemlineata* (Coleoptera: Chrysomelidae)) evaluated on tobacco (*Nicotiana tabacum* (Solanales: Solanaceae)) and potato (*Solanum tuberosum* (Solanales: Solanaceae)) plants, respectively. The small-bodied insects were European corn borer (*Ostrinia nubilalis* (Lepidoptera: Crambidae)) and Northern corn rootworm (*Diabrotica barberi* Smith & Lawrence (Coleoptera: Chrysomelidae)), on corn plants (*Zea mays* (Poales: Poaceae)).

To test the hypothesis that a contact microphone clipped onto a plant could detect insect herbivory, we implemented a randomized paired study design, wherein two plants were simultaneously recorded with and without insect(s) was used. Recordings occurred at the University of Wisconsin-Madison (Room 537E, 1630 Linden Dr, Madison, WI 53706) within a sound-dampened room linked with moving 24 137cm x 188cm textile moving blankets (Nova Microdermabrasion, China) affixed to all walls to dampen noise, and continuously lit with eight 25V 5600 K lights (GE Ecolux, Boston, Massachusetts USA) throughout the measurement periods. A contact microphone was clipped to the base of each of the paired plants five cm above the soil (Figure 1B). Each experiment was run for one hour, with one of the plants in each pair containing an insect, or insects, in the case of Northern corn rootworm. For the two species of large-bodied insects (Russell Labs colony, 1630 Linden Dr., Madison WI 53706), individuals were randomly placed on one of the plants. For the small-bodies insects, third-instar European corn borers (Benzon Research, 7 Kuhn Drive, Carlisle, PA 17015) were individually placed on a subset of pest-free plants. Two thousand Northern corn rootworm eggs were injected into the soil, resulting in an estimated one to seven successfully hatched larvae, confirmed using a Berlese trap. For each plant containing small-bodied insect(s), the plant was randomly assigned placements for the experiment. Plants were left unobserved for ten minutes before the start of the study for insect acclimation. The tobacco plants were placed on a granite slab for noise reduction. Potato and corn were placed on 12mm x 300mm x 200 mm neoprene foam mats (RHBLME, China) to limit the recording of building vibrations and thus lower signal noise.

To ensure that the small-bodied insects were monitored during periods of active feeding, one 24-hour evaluation of circadian feeding activity was performed. This observation was only performed with the small-bodied insects, as feeding was visually observable in the large-bodied insects during the experimental period.

A field study evaluated the sensor’s insect eavesdropping capability *in situ*. On 06/13/2023, in a field (Arlington Research Station, University of Wisconsin-Madison 43°18’54.7”N 89°20’24.4”W) managed for high corn rootworm (*Diabrotica* spp.) larval density using continuous corn planting, a sensor was set up with two contact microphones clipped to two corn plants. Plants either were untreated or were planted alongside an in-furrow insecticide (Capture LFC 8.5 fl. oz. / acre). Twice a day, 9:00 -11:00 and 15:00 – 17:00, the contact microphones recorded contact microphones. Additionally, plants were visually inspected for above-ground herbivory, and at roots were dug on 7/15/2023 to confirm and quantify corn rootworm damage.

Insect presence and absence were evaluated by comparing the number of peaks, where each peak is hypothesized to represent insect interaction with a plant. Peaks were detected using the ‘librosa.onset.onset_detect’ from the ‘librosa’ library (https://zenodo.org/records/11192913).

In initial exploratory analysis using a Fourier transform of the recorded audio signal, a 60Hz signal was found to be present across all recordings. This is expected to be caused by electrical interference from the AC power source used by the Raspberry Pi. An IIR notch digital filter was therefore applied to the signal using scipy.signal.iirnotch(1) with a quality factor of 30 to remove 60Hz frequencies, as well as the first 7 harmonic frequencies. This substantially reduced the baseline noise in the data.

To separate periods of potential insect activity from background noise, the standard deviation of the signal was calculated, and a threshold set at 20-times the standard deviation independently for each trial. Values above this threshold were separated into distinct activity bouts by merging high-amplitude signals occurring within 0.01 seconds of one another. The resulting regions are then dilated to include the 100 frames (∼0.000045 seconds) prior and 200 frames (∼0.00009 seconds) following to ensure that the region of activity was not truncated.

## Results

The sensor was successfully able to record feeding activity from all four evaluated insect species, as determined by a disproportionately greater number of peaks extracted from plants containing insects, compared to the insect-free control plants. Specifically, the sensor detected tobacco hornworms, Colorado potato beetles, European corn borers, and Northern corn rootworms feeding in a laboratory setting (Figure 2A-D).

**Figure 2.**
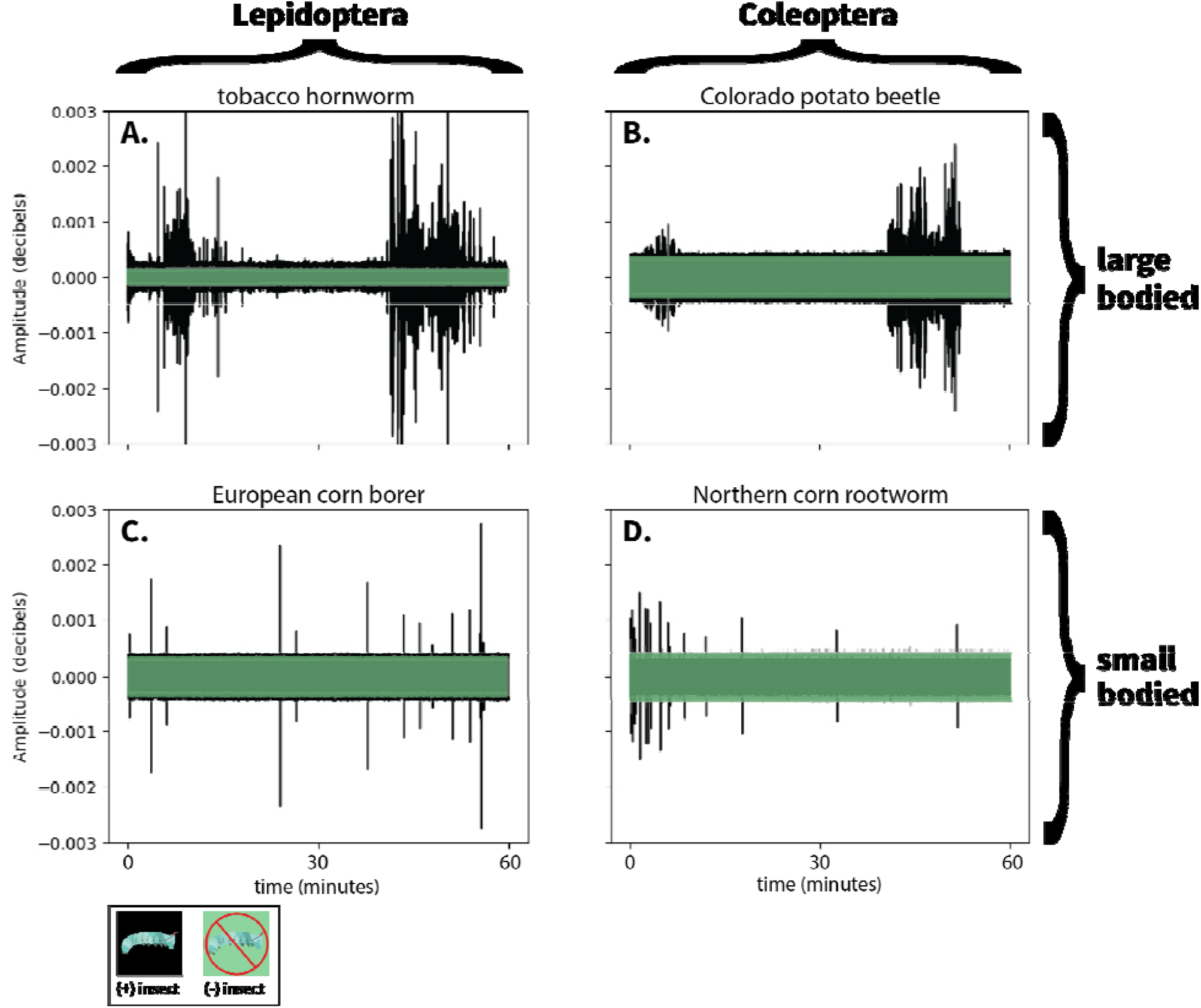
Example signals from one hour of recordings for plants with insects (in black) and without insects (in gray) extracted from contact microphones clipped onto **A**. tobacco plants with and without tobacco hornworm (upper left), **B**. potato plants with and without Colorado potato beetle (upper right), **C**. corn plants with European corn borer (lower left), and **D**. corn plants with Northern corn rootworm (lower right).

**Figure 3.**
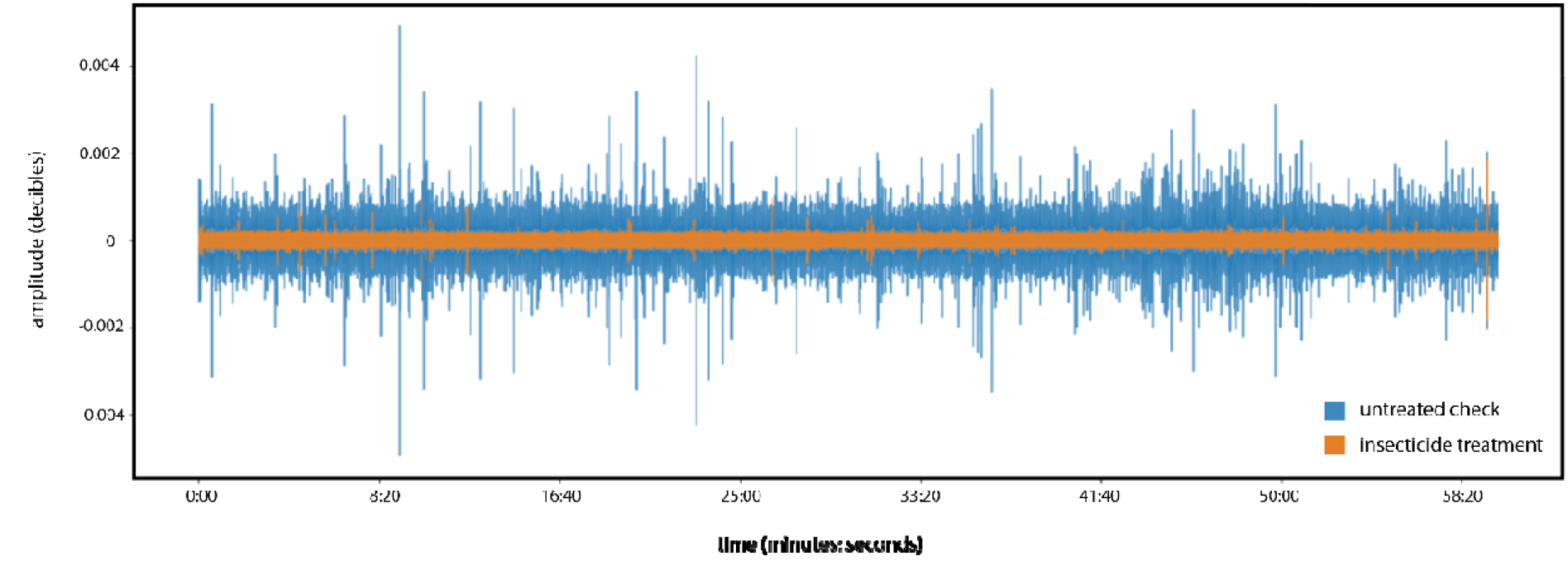
Amplitude of signal recorded from the sensor clipped onto corn plants in a continuous corn field. Sensors were clipped to corn plants treated either with an insecticide in-furrow applied insecticide (orange) or left untreated (blue).

Differences in feeding sounds from corn rootworm species (*Diabrotica* spp. (Coleoptera: Chrysomelidae)) were observed in an agricultural corn field. Specifically, within an hour-long recording, there were greater numbers of sounds above the 20x baseline threshold for plants with no insecticide (1888 sounds identified thought to be insect feeding), than the plant that was treated with an in-furrow insecticide application (12 sounds identified as feeding). Moreover, the amplitudes of the control plant’s sounds were greater than the insect feeding sounds on the treatment plant.

## Discussion

Cost-effective insect sensors hold great potential for elucidating plant-insect interactions and improving agricultural decision-making. Here, we demonstrate that low-cost clip-on contact microphones ($67 per microphone) can detect insect herbivory in both a laboratory and field setting for large and small-bodied species. This type of sensor allows for unprecedented real-time observation of below ground and boring insect behavior. Moreover, the device and method allow for continuous monitoring, enabling observations that were previously impossible using destructive sampling (e.g., changes in herbivore activity over time within plants or below ground).

The use of a cost-effective contact microphones is the most recent advancement in over a century of recording audio and vibrations for studying insect behavior (Mankin et al., n.d.). In 1964, the electroactograph was described as a technique for measuring activity rhythms in lepidopteran defoliators. Ten years later, a sound cartridge and recording tape recorded mating signal substrate vibrations of the brown planthopper. Just two years following, a condenser microphone in a laboratory was used to elucidate the mating behavior of two taxa in the *Macrosteles fascifrons* species complex. Condenser microphones have extremely high signal to noise ratios, enabling this observation. Laser vibrometers were applied in 1982 to measuring pre-mating substrate communication of multiple hemipteran species, and in 1984 accelerometers were applied. The benefits and limitations of each of these methodologies are explored in ‘Vibrational Communication in Insects’. Two major differences exist between these methods and the one described here: (1) the vast majority of these recordings were performed in a controlled laboratory environment, and (2) methods often do not take advantage of signal processing algorithms for amplifying the noise to signal ratio. Moreover, the detection of soil insects with sensors primarily occurs using soil probes, rather than direct detection of insect’s interactions with plants.

The approaches described here holds great potential for understanding insect herbivory behavior and has widespread potential applications in agriculture. In agriculture, this type of sensor could help a grower gain valuable insights into intervention efficacy and potentially assist with timing of future interventions. Cost often excludes the use of insect sensors at scale as well as their initial adoption. When sensors are cost effective, they can be run in a network, expanding the power of the sensors to allow for interpolation (Potamitis et al., 2017). This is likely the best use case in an agricultural setting, with data interpolated between sensors.

Our approach could also be used across diverse insect and plant taxa: Much research on automated monitoring has focused on pollinators due to the importance the important ecosystem functions they provide (Bjerge Id et al., 2023; Pegoraro et al., 2020); contact microphones could likely detect floral visits of pollinators, especially bee-mediated buzz polination, to flowers for spatially explicit evaluations their pollinator presence, movement, and identification. Moreover, the approached developed here is robust to plant species and could be used to monitor insect visits to a range of plants across an arena or a field with applications in pesticide monitoring and environmental monitoring

While we demonstrate here the potential for the application of substrate-borne acoustic signals to detect insect herbivores, real-world application in agricultural ecosystems will require refined algorithms and an expanded toolkit for detecting and identifying insects. For example, algorithms for identifying insect vibrations from other environmental noise – as developed here – will need to be tested and validated across a broader range of systems. Next, for many applications, algorithms enabling identification of insect species or species complexes will be required. The problem space – or list of possible insect species – is somewhat limited by the plant or crop and its insect community, easing insect identification. Finally, a relationship between the density of insect species and the feeding rates needs to be established. Preliminary work suggests the application of deep learning to these questions may be a successful avenue for future exploration.

While the results here provide a ‘proof-of-principle’ for low-cost acoustic monitoring, the sensor hardware presented here has important limitations. First, the sensor’s observation is limited to a single plant per microphone by its need for each microphone to be clipped onto a plant. A single plant observation requires replication for it to be representative in a population, especially when bringing these sensors to agricultural systems. Currently, the sensor requires the physical removal of memory storage, which may be challenging in field conditions. The device could be further improved by the ability to transmit data remotely via Bluetooth, LARA, or WIFI. This would likely require edge computing, or the extraction of likely insect events on the sensor. Wiring poses challenges for field use, as it is exposed to the elements, and is at risk of damage.

In conclusion, this sensor allows for eavesdropping on plant-insect feeding interactions. The low cost of the sensor likely will ease the transition to adoption for a range of applications, from basic science research to applied agricultural work. While this sensor is not the first audio-vibration measurement device, its ability to record soil dwelling insects’ feeding appears to be unique. With the application of existing algorithms, authors foresee a wide range of future potential use cases.

## Acknowledgments

We would like to acknowledge Dr. Russell Groves for facilitating the potato work, Grace Melone and Ann M for growing the insects, Alex Arovas for reviewing the …

Additionally, we would like to acknowledge x for providing seeds for the project. FMC for pesticide donations; Russ’s field crew for planting; WARF Accelerator grant…

## Author Contributions

EB conceptualized the use case of contact microphone application for insect monitoring. DM, LS, VA, and JC assembled the sensor and code. KG and EB conceived of experimental methods. DM, KG, and EB conducted experiments. DM and EB generated results. EB led writing of the manuscript. DM, KG, JC, and EB edited the manuscript. All authors contributed critically to the drafts and gave final approval for publication.

## Data archive intension

https://github.com/wiscbicklab/InsectEavesdropperMethodsPaper

